# Filamentation in *Candida albicans* is modulated by adaptive translation of farnesol signalling genes

**DOI:** 10.1101/2021.01.20.427544

**Authors:** Carla Oliveira, Ana Rita Guimarães, Inês Correia, Inês Sousa, Ana Poim, Sílvia M. Rocha, Gabriela Moura, Manuel A. S. Santos, Ana Rita Bezerra

## Abstract

The complex biology of the human pathogen *Candida albicans* is reflected in its remarkable ability to proliferate in numerous body sites, adapt to drastic changes in the environment, form various types of colonies and grow in yeast, pseudo-hyphal and hyphal forms. Much has been learnt in recent years about the relevance of this phenotypic plasticity, but the mechanisms that support it are still not fully understood. We have demonstrated that atypical translation of the CUG codon is a source of unexpected morphological diversity. The CUG codon is translated as both leucine (Leu) (~3%) and serine (Ser) (~97%) in normal growth conditions, but Ser/Leu levels change in response to stress. Remarkably, recombinant *C. albicans* strains incorporating between 20% and 99% of Leu at CUG sites display a diverse array of phenotypes and produce colonies of variable morphology containing a mixture of yeast, pseudohyphal and hyphal cells. In this work we investigate the role of the CUG codon in the yeast-hypha transition. Our data show that increasing incorporation levels of Leu at CUG sites trigger hyphal initiation under non-inducing conditions by reducing farnesol production, and increasing the degradation of the Nrg1 hyphal repressor. We propose that dual CUG Ser/Leu translation triggers filamentation via the Nrg1 pathway.

**Importance:** The unique translation of the CUG codon as both Ser (~97%) and Leu (~3%) plays a key role in the production of high genomic and phenotypic diversity in *C. albicans*. The molecular mechanisms that support such diversity are poorly understood. Here, we show that increased Leu incorporation at CUG sites induce hyphae formation in media where *C. albicans* normally grows in the yeast form. The data show that increasing Leu at CUG sites triggers the degradation of the hyphal repressor Nrg1, allowing for full expression of hyphal genes. Since filamentation is important for invasion of host tissues, this work shows how the atypical translation of a single codon may play a critical role in the virulence of all fungi of the CTG clade.

## Introduction

*C. albicans* is able to undergo reversible morphological changes from yeast (single oval cells) to filaments (elongated cells) in response to environmental cues (1). Filamentous cells, including hyphal and pseudohyphal morphologies, are linked to the establishment of biofilms, escape from phagocytes and invasion of host endothelial and epithelial tissues (2, 3). Yeast cells are more suited for dissemination through the bloodstream (4), but both morphologies are essential for full virulence as strains locked in either one of them have attenuated virulence in a mouse model of systemic candidiasis (4–6).

Hyphal development is regulated by a complex interplay between external and internal factors. External factors include environmental cues that reflect the diversity of the microenvironments encountered in the host, such as high temperature (37°C), serum, neutral pH, low levels of oxygen, high levels of CO_2_, N-acetyl-d-glucosamine (GlcNAc), and poor nutrition conditions (7–9). Numerous molecules have also been implicated in this process, specifically fatty acids, peptides, cell cycle inhibitors and quorum-sensing molecules (QSMs), such as farnesol (10). Most of these signals induce filamentation through the activation of adenylyl cyclase Cyr1 or through the small GTPase Ras1 which in turn activates Cyr1 (8, 11). The target of Cyr1 is the cyclic AMP-protein kinase A (cAMP-PKA) cascade through the activation of the PKA catalytic subunits Tpk1 and Tpk2. These can phosphorylate target proteins or bind to promoter regions of target genes to modulate their transcription (12, 13). One of these targets is Nrg1, a key repressor of the hyphal transcriptional program (8, 14–16), which acts on the promoters of hypha-specific genes, such as *ECE1* and *HWP1*, to repress their expression during yeast growth. Thus, hyphal development involves two stages: 1) repression of Nrg1 expression during initiation; 2) removal of Nrg1 during hyphal maintenance by blocking its capacity to bind to promoters of hypha-specific genes (16, 17). Two independent pathways are involved in clearing Nrg1 during initiation, namely transcriptional downregulation of Nrg1 through activation of the PKA cascade, and Nrg1 degradation through the reduction of farnesol levels (16–18).

Farnesol is a byproduct of the ergosterol biosynthesis pathway (19) that supresses the Ubr1-mediated degradation of Cup9, a transcription factor that represses the expression of *SOK1*. The repression of Sok1 expression blocks Nrg1 degradation and hyphal initiation. Hence, removal of farnesol triggers Cup9 degradation, resulting in Sok1 expression and Nrg1 degradation leading to hyphal initiation (16–18).

We have discovered that *C. albicans* normally translates CUG codons mainly as Serine rather than the typical Leucine found in other organisms. However, exposure to oxidative and pH stress leads to an increase in Leu incorporation levels from 3% up to 5%, producing functional statistical proteins whose sequence is related to the sequence of the wild-type (wt) peptide by some statistical measure – with the concurrent impact on the pathogen’s biology (20). Recombinant *C. albicans* strains incorporating between 20% and 99% of Leu at CUG sites are viable and display remarkable phenotypic variability, including colonies with highly variable morphologies. These cells grow on agar plates as wrinkled colonies due to the high proportion of pseudophyphal and hyphal cells within the colonies; smooth colonies of wt strains contain mainly yeast-like cells (21, 22).

Considering that 10% of CUG codons are located in conserved positions of proteins involved in signaling pathways that regulate filamentous growth (23), we hypothesized that variable Ser/Leu incorporation at CUG positions may destabilize protein structure or spawn new protein functionalities that may modulate hyphal growth. To test this hypothesis, we used *C. albicans* strains that incorporate Leu at CUG sites at high level (21). Germ-tube (GT) assays performed in non-inducing conditions showed that approximately 50% of cells of these strains had germinating characteristics, while only 1.4% of control cells showed such features. Their metabolomic profiles showed lower concentrations of the quorum sensing metabolite farnesol, which is involved in the degradation of the filamentation repressor Nrg1. How increased Leu incorporation at CUG sites decreases the production of farnesol is not clear, but this study shows that atypical CUG translation triggers filamentation by altering quorum-sensing signaling in *C. albicans*.

## Results

### Leu-CUG translation regulates filamentation under non-inducing conditions

We have constructed a series of *C. albicans* strains that incorporate 20% (T1), 50% (T1KO1), 80% (T2KO1) and 98% (T2KO2) of Leu at CUG sites (see Table S1) (21). Previous phenotypic studies indicate that increasing levels of Leu incorporation at CUG sites decreases growth rate but generates morphological diversity. Such strains produced colonies with irregular-wrinkled and jagged morphologies composed of a mixture of yeast, pseudohyphal, and hyphal cells (21, 22). To further characterize the effects of Leu incorporation at CUGs (Leu-CUG) on morphology, we used germ-tube (GT) formation assays to measure filamentation under various growth conditions (17, 24) . In non-inducing conditions (YPD at 30°C), increased Leu-CUG had a profound impact on filamentation as T1, T1KO1, T2KO1 and T2KO2 cells produced between 25-45% of germ tubes within 5 hours of growth while less than 2% of wt cells filamented in the same conditions (Fig. 1A). On the other hand, upon induction of GT formation at 37°C and in serum, control and T1, T1KO1, T2KO1 and T2KO2 cells produced similar levels of germinating cells (~80%) (Fig. 1A). As expected, wt cells grown at 37°C and in serum showed normal germ tube formation after 5 hours of growth and sustained hyphal growth after 24 hours (Fig. 1B). The same phenotype was observed for T1 cells (20% Leu) (Fig. 1C). In contrast to the wt population that consistently grew as round yeast cells at 30°C (Fig.1B), T1 cells showed the same pattern of sustained hyphal growth in both inducing and non-inducing conditions (Fig. 1C). Therefore, hyphal development is not affected by Ser/Leu ambiguity at CUG positions in hyphal inducing conditions, but high levels of CUG- Leu translation contribute significantly to the initiation of filamentation in non-inducing conditions.

**Fig 1.**
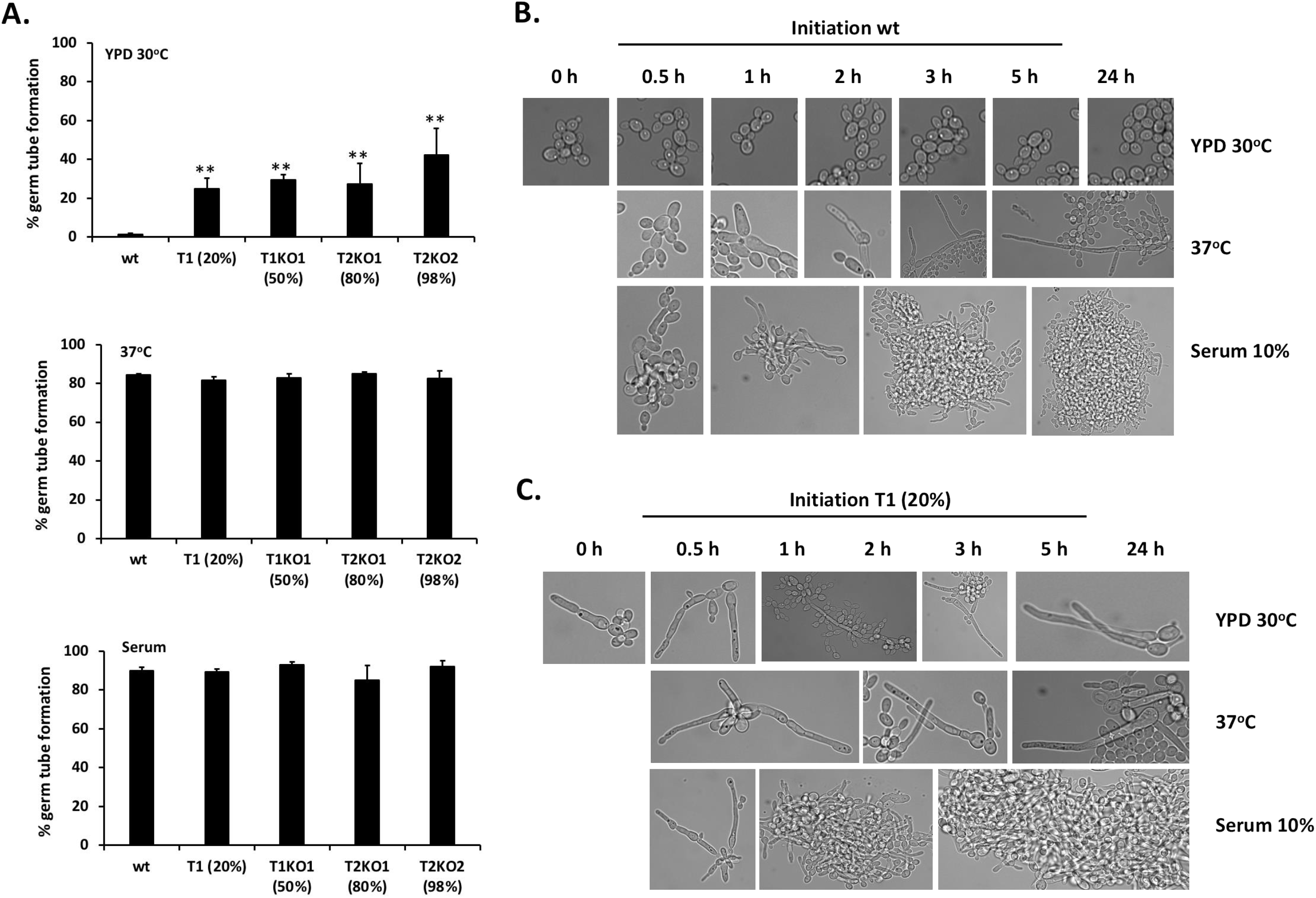
Leu-CUG translation regulates filamentation under non-inducing conditions. **A.** Filament induction of wild type and high-level Leu-CUG translating strains (T1, T1KO1, T2KO1 and T2KO2) incubated for 5 hours under non-inducing conditions (YPD 30°C) and under hypha-inducing conditions (RPMI 370C and RPMI with 10% serum). **B.** Morphology of wild type cells grown under 30°C, 37°C and with serum. **C.** Morphology of the high-level Leu-CUG translating T1 strain cells grown under 30°C, 37°C and with serum. The microscopic pictures reflect a representative example of the observed phenotype for each strain and condition. Scale bar: 10 μm. Data are presented as mean and SD from three independent biological replicates. Asterisks indicate statistically significant differences (\**P* < 0.05; ** *P* < 0.01; *** *P* < 0.001) determined by Student t test.

### Incorporation of Leu at CUG sites downregulates farnesol production

Since our previous genomic studies did not explain the filamentation phenotype of T1, T1KO1, T2KO1 and T2KO2 strains (21), metabolomic changes associated with the differential morphogenesis of these strains in non-inducing conditions were assessed using comprehensive two dimensional gas chromatography–mass spectrometry, with time of flight analyser (GC×GC–ToFMS). We detected over 1000 GCxGC instrumental features that were identified using mass spectral libraries (see Materials and Methods for full details). These compounds were distributed over a wide range of chemical families: acids, alcohols, aldehydes, hydrocarbons, esters, ketones, monoterpenic and sesquiterpenic compounds, norisoprenoids, phenols, and sulphur compounds. Multivariate statistical analyses of the data allowed us to identify 16 key metabolites that differed in concentration between the strains that incorporate Leu at high level and the control cells. These metabolites and their putative biological function and changes in levels of expression are shown in Figure 2A (see also Table S2). Five of them, including 2,4-dimethyl-1-heptene, 4-methyl-octane, 2-propenal, 2,6-dimethyl-4-heptanone, and dihydromyrcenol do not have a functional pathway attributed in current databases and literature. Using the KEGG database, the remaining 11 could be associated with metabolic pathways such as gluconeogenesis, pyruvate metabolism, energy production and Leu degradation (25). These metabolites have been previously implicated in the morphogenetic process of *C. albicans* during engulfment by macrophages or when cells are cultivated in hyphae-inducing media (26, 27).

**Fig 2.**
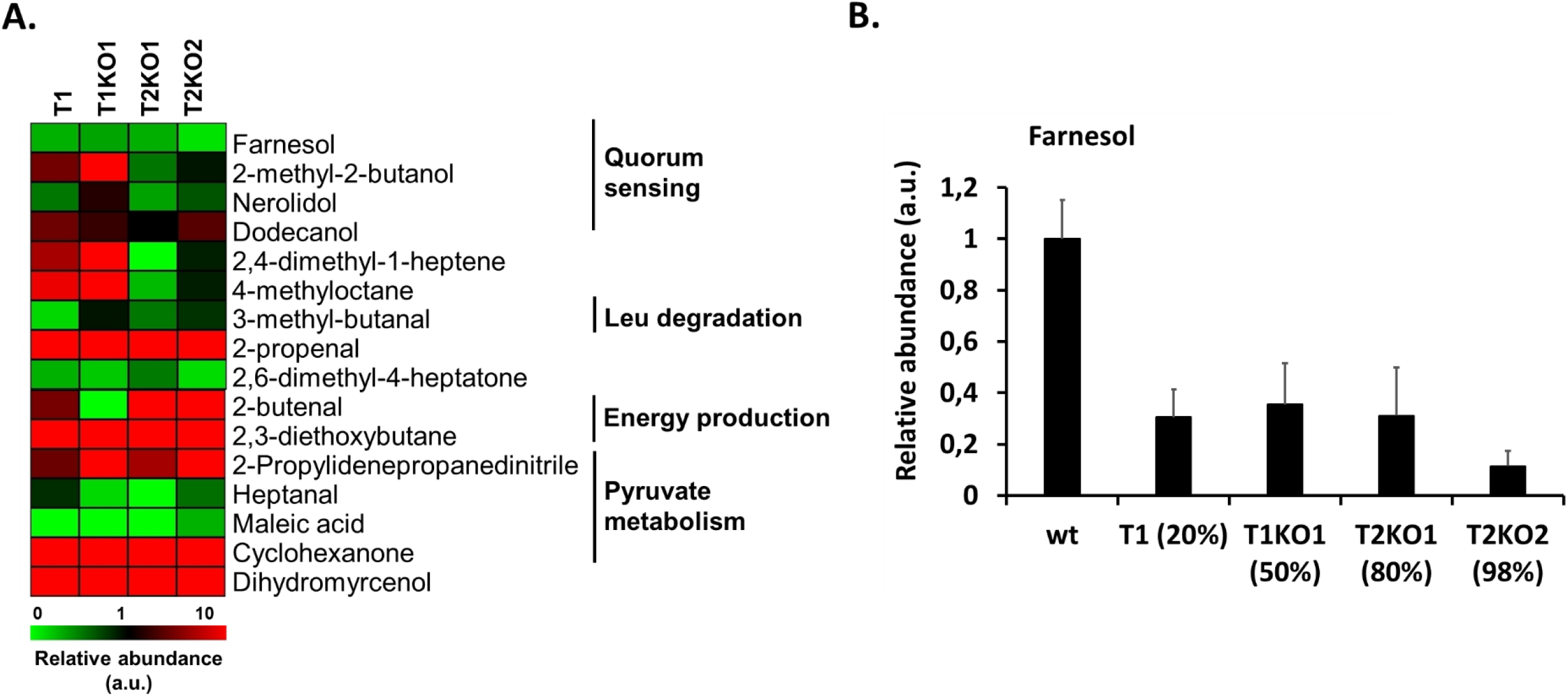
Metabolite profiles of high-level Leu-CUG incorporating cells using GC×GC-ToFMS. **A.** Relative abundances of metabolites detected by GC×GC-ToFMS in the strains that incorporate Leu at high level T1, T1KO1, T2KO1 and T2KO2 when compared to wt. Black squares represent a concentration that is indistinguishable from the wt strain. Green/red squares represent lower/higher metabolite abundance of the test strain relative to wt, respectively. Only metabolites with statistically different abundances in at least one of the high-level Leu-CUG translating strains are shown. **B.** Farnesol abundance was normalized to the abundance measured in wt. Its abundance was lower in strains that incorporate Leu at high level. Data are presented as mean and SD from three independent biological replicates. Asterisks indicate statistically significant differences (\**P* < 0.05; ** *P*< 0.01; *** *P* < 0.001) determined by Student t test.

Four metabolites were detected in more abundance in the T1, T1KO1, T2KO1 and T2KO2 strains, namely 5-methyl-2-(1-methylethyl)-cyclohexanone and 2-propylidenepropanedinitrile, which participate in pyruvate metabolism and, 2,3-diethoxybutane and 1-dodecanol, which are energy production and quorum sensing metabolites, respectively (Fig. 2A). Conversely, 4 metabolites had lower abundance in all strains, namely heptanal, maleic acid and 3-methyl-butanal, which are involved in the pyruvate and Leu metabolic processes, and farnesol, a well-known quorum sensing molecule (19). Strains T1, T1KO1 and T2KO1 produced approximately 70% less farnesol than the control and strain T2KO2 downregulated farnesol by 90% (Fig. 2B*). C. albicans* produces high amounts of farnesol as a byproduct of the ergosterol biosynthesis pathway and the most prominent role of this molecule is the inhibition of the yeast-to-hypha transition (17, 18, 28). Its lower abundance in all strains tested (Fig. 2B) was correlated with growth in the filamentous form and production of highly wrinkled colonies.

To validate our data, 100 μM of farnesol was added to YPD cultures of T1, T1KO1, T2KO1 and T2KO2 strains. Germ tube formation was reduced from 25-45% to approximately 2%, i.e., to nearly wt levels (Fig. 3).

**Fig 3.**
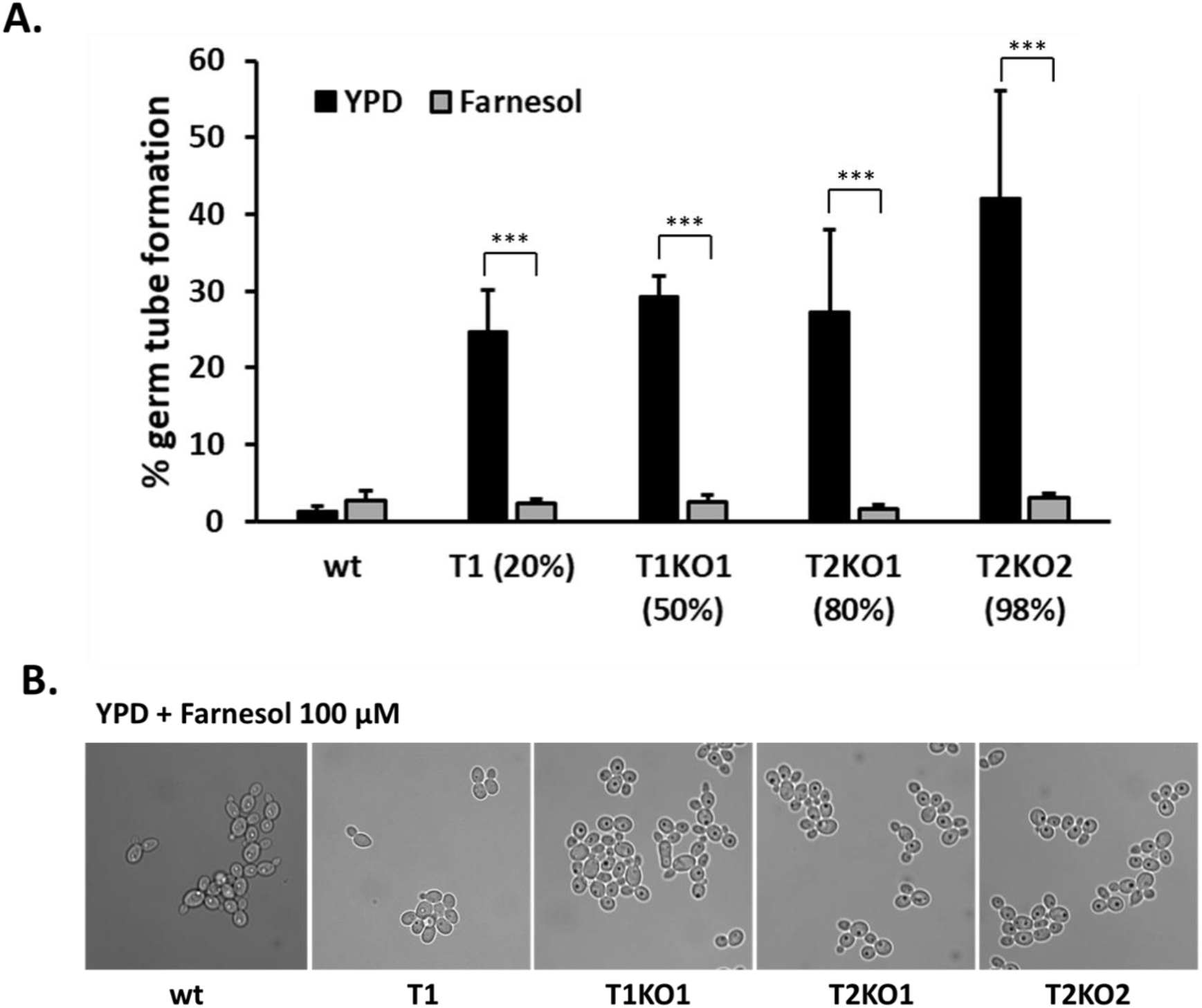
The effect of exogenously added farnesol on germ tube formation of wt and high-level Leu-CUG translating strains. **A.** Filament induction of strains incubated for 5 hours under non-inducing conditions (YPD 30°C – black bars) and in the presence of farnesol (FOH 100 μM – grey bars). B. Morphology of cells grown with farnesol. The microscopic pictures reflect a representative example of the observed phenotype for each strain. Scale bar: 10 μm. Data are presented as mean and SD from three independent biological replicates. Asterisks indicate statistically significant differences (\**P* < 0.05; \*\**P* < 0.01; \*\*\**P* < 0.001) determined by Student t test.

Since farnesol blocks the degradation of the repressor of the hyphal initiation pathway (Nrg1) (17, 18), these results suggested that Leu-CUG translation may trigger filamentation by decreasing farnesol production and consequently prompting Nrg1 degradation.

### Leu-CUG translation increases degradation of Nrg1

A previous study has shown that releasing cells from farnesol inhibition activated the degradation of the Cup9 transcriptional repressor, allowing for the expression of the Sok1 kinase and subsequent Nrg1 protein degradation, which in turn derepresses hyphal initiation (18). Considering this, we asked whether the Nrg1 protein was degraded in the high-level Leu-CUG incorporating cells, due to low farnesol levels. Wt and T1 (20% Leu) cells, expressing recombinant Nrg1 fused to a polyhistidine tag (His_6_-Nrg1) were inoculated from overnight cultures into fresh YPD media, at 30°C with and without addition of 100 μM of farnesol. Immediately after inoculation into non-inducing medium (0h), Nrg1 protein levels decreased sharply in T1 cells relative to control cells; and continued to decrease after 5 hours of growth (Fig. 4A), indicating that Leu-CUG translation affected Nrg1 protein stability. Adding farnesol to T1 cells blocked Nrg1 degradation and cells recovered Nrg1 to wt levels (Fig. 4B), validating the hypothesis that increased Leu-CUG translation activates degradation of the hyphal repressor through the quorum sensing metabolite farnesol.

**Fig 4.**
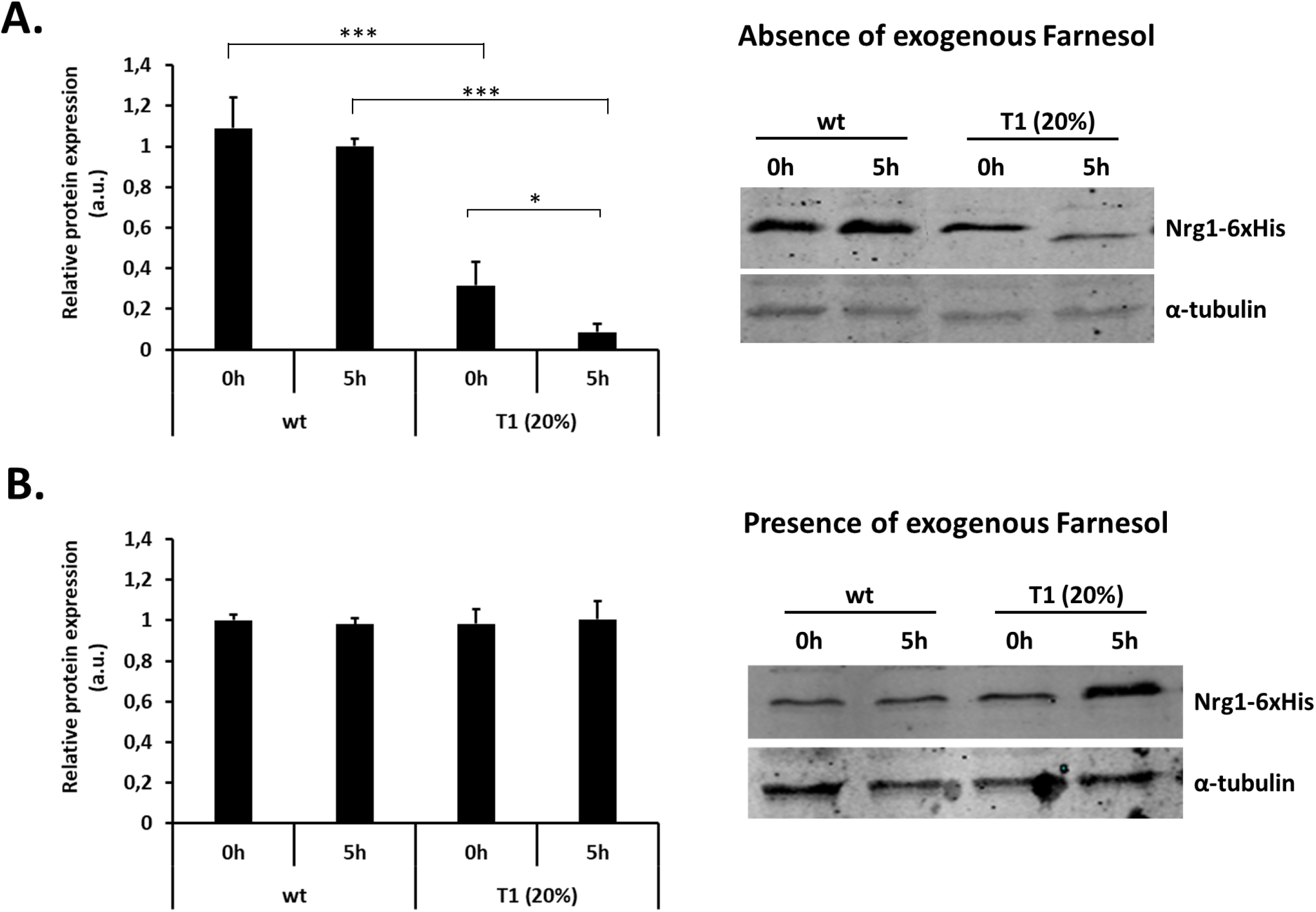
Leu-CUG translation increases degradation of Nrg1. **A.** Nrg1 protein levels in wt (T0) and high-level Leu-CUG translating cells (T1-20%) grown in non-inducing conditions. Cells expressing recombinant Nrg1 fused with a His_6_-tag were subcultured to log phase (5h) in YPD. **B.** Nrg1 protein levels in cells subcultured to log phase in YPD plus farnesol 100 μM. Total protein was resolved by SDS-PAGE, and the blot was hybridized with anti-His_6_-tag to detect Nrg1 and anti-α-tubulin to monitor α-tubulin as a loading control. Data are presented as mean and SD from three independent biological replicates. Asterisks indicate statistically significant differences (\**P* < 0.05; ** *P* < 0.01; *** *P* < 0.001) determined by Student t test.

Considering that most proteins of the farnesol-sensing hyphal initiation pathway contain amino acid residues encoded by the CUG codon, we hypothesized that degradation of the repressor in T1, T1KO1, T2KO1 and T2KO2 could be caused by direct destabilization of protein structure/function by Leu incorporation at Ser CUG sites. To test this hypothesis, we evaluated whether the Ser-for-Leu substitution had direct impact on protein structure, using an *in silico* approach based on the PROVEAN (Protein Variation Effect Analyzer) algorithm (29). PROVEAN searches homologs of a given protein in the NCBI database using BLAST and clusters them using the CD-HIT program. Using the selected homologs, PROVEAN scores are computed for each of the variants provided (see Materials and methods). The impact of the Ser-for-Leu substitution at CUG sites was predicted to be neutral in all proteins of the farnesol-sensing hyphal initiation pathway, including Cup9, Sok1 and Nrg1 (Table S3), suggesting that Nrg1 protein degradation was indeed a consequence of the absence of farnesol, rather than a direct consequence of the Ser-for-Leu substitution at the CUG sites of those proteins.

To further validate that Leu incorporation at the Nrg1 CUG-207 site does not destabilize the Nrg1 protein, we constructed 3 new *C. albicans* strains that expressed the different CUG-site isoforms of Nrg1. To do so, the CUG-207 site of the *NRG1* gene was mutated to both Leu-UUA and Ser-UCU codons (Table S1). Therefore, the strain T0_CUG207 expressed the ambiguous (Ser/Leu) wt Nrg1 protein, while strains T0_UUA207 and T0_UCU207 expressed mutant Nrg1 with only Leu or Ser at position 207, respectively (Fig. 5A). Western blot analysis showed no difference in Nrg1 protein levels between the different isoforms, confirming that the Ser-for-Leu replacement at the CUG-207 site did not affect protein stability (Fig. 5B). Furthermore, the function of the protein was not affected because cells expressing the Ser-207 or Leu-207 isoforms of Nrg1 grew in the yeast form, which would not be the case if the hyphal repressor was not functional, as demonstrated by the filamentous growth of Δ*nrg1*/ Δ*nrg1* mutant cells (Fig. 5C).

**Fig 5.**
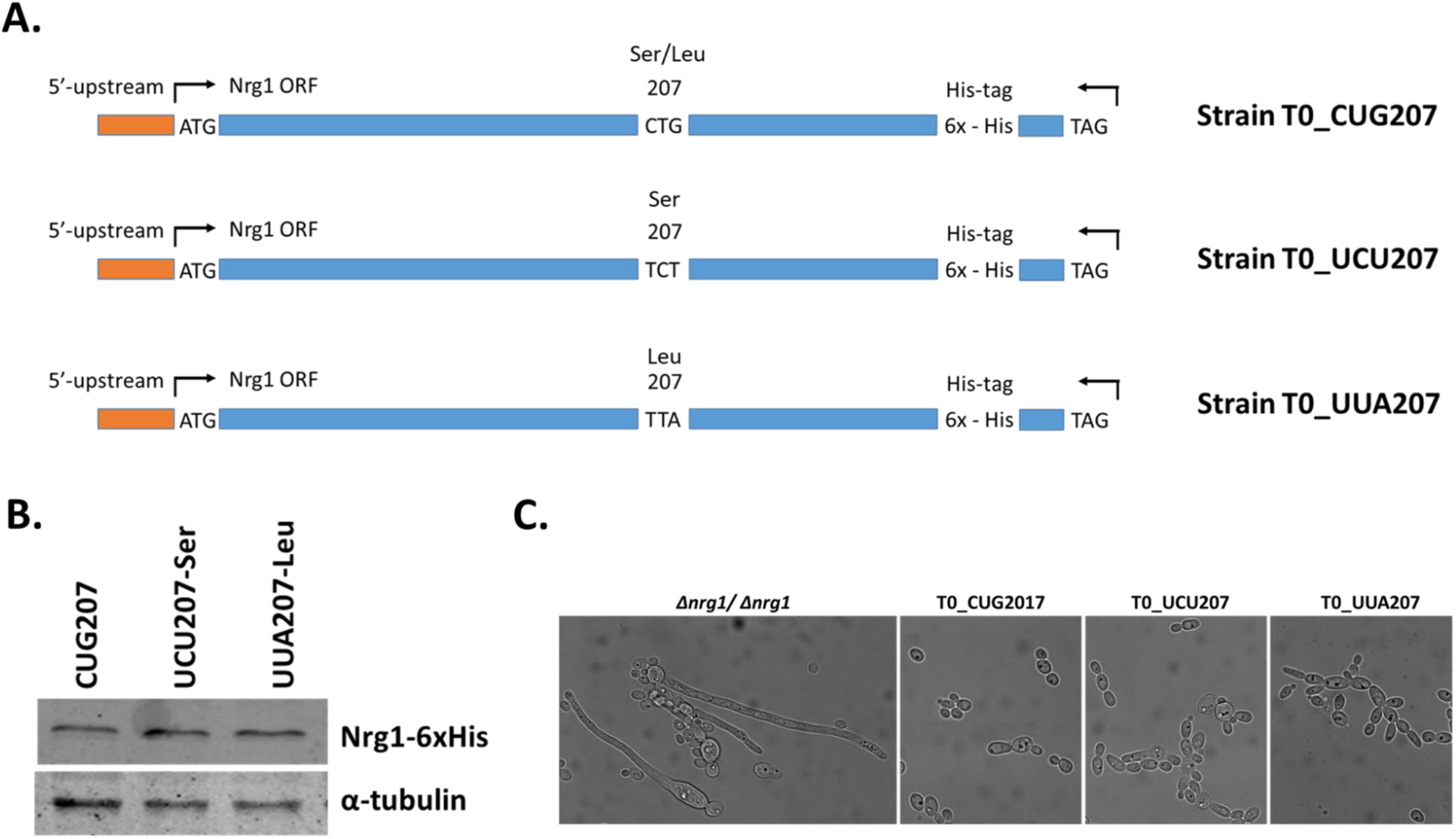
Leu-CUG translation does not directly destabilize Nrg1 (CUG-207). **A.** Engineered strains of *C. albicans* expressing one of the two Nrg1 isoforms only. The *NRG1* gene encoding a CUG codon at position 207 was substituted by a *NRG1* gene with a Leu UUA codon and a Ser UCU codon at position 207. In all cases *NRG1* is fused with His_6_-tag **B.** Western blot analysis showing that *C. albicans* cells expressing the UUA_Leu207 and UCU_Ser207 isoforms produce the same amount of Nrg1. **C.** Morphology of cells expressing the UUA_Leu207 and UCU_Ser207 isoforms, grown under non-inducing conditions. The microscopic pictures reflect a representative example of the observed phenotype for each strain. Scale bar: 10 μm.

### Leu-CUG translation did not alter transcriptional regulation of Nrg1

Since farnesol is also involved in the transcriptional downregulation of Nrg1 (16, 18), we used DNA microarrays to clarify if the hyphal initiation pathway was downregulated. The experiment focused on the T1 strain (20% Leu) whose genome was previously sequenced and did not display large-scale genomic rearrangements (21). Only genes with a *P value* inferior to 0.01 were considered in the analysis (Dataset S1). There was no transcriptional downregulation of *NRG1* or deregulation of other genes involved in this pathway, specifically *CYR1* and *TPK2*. On the other hand, one gene that belongs to the Nrg1 protein degradation pathway (*SOK1*) was upregulated (Table 1), supporting a previous study where SOK1 expression was activated upon release from farnesol (18).

**Table 1.** Gene expression in the high level Leu-CUG translating strain T1 as revealed by transcript profiling

| Gene | Functional description | Log FC | *p*-value |
| --- | --- | --- | --- |
| **Genes involved in downregulation of Nrg1 during hyphal initiation (21)** |  |  |  |
| **CYR1** | class III adenylyl cyclase; involved in regulation of filamentation | -0,123 | 0,406 |
| **TPK2** | protein kinase; involved in regulation of filamentation | -0,650 | 0,061 |
| **CUP9** | transcription factor; represses SOK1 expression in response to farnesol inhibition | -0,385 | 0,012 |
| **SOK1** | protein kinase required for degradation of Nrg1p | 1,683 | 1,19E-06 |
| **NRG1** | transcription factor/repressor; regulates hyphal gene induction | 0,222 | 0,390 |
| **Hyphal-specific genes controlled by Nrg1 (16-17)** |  |  |  |
| **ALS3** | cell wall adhesin; epithelial adhesion, endothelial invasion | 0,087 | 0,796 |
| **CEK1** | ERK-family protein kinase; required for wild-type yeast-hypha switch | 0,846 | 7,10E-04 |
| **ECE1** | candidalysin; hypha-specific protein | 1,045 | 7,80E-06 |
| **HGC1** | hypha-specific G1 cyclin-related protein involved in regulation of morphogenesis | 0,296 | 0,457 |
| **HWP1** | hyphal cell wall protein | 1,646 | 7,61E-03 |

Fold regulation for a specific gene = (normalized mean signal in high Leu-CUG translating / normalized mean signal in wt)
Genes whose expression in significantly altered (*p*-value<0.01) are highlighted

Since Nrg1 represses a set of hyphal-specific genes (6, 14, 15), we were expecting that these genes would be induced as a direct result of its downregulation in the T1 cells. The transcriptome data confirmed that expression of *CEK1*, *ECE1* and *HWP1* was upregulated (Table 1), though the fold-change variation relative to the wt was relatively small.

The transcript profiling dataset did not allow us to identify alterations in known signaling pathways associated with farnesol production, particularly in the ergosterol biosynthetic pathway.

## Discussion

Environmental triggers such as low temperature, CO2, amino acids and glucose, regulate hypha formation in *C. albicans* through the cyclic AMP (cAMP)–protein kinase A (PKA) signalling pathway. Most signals enhance filamentation by acting on the adenylate cyclase Cyr1 pathways, while others act through the transcription factor Efg1 to activate hypha-specific cyclins that form complexes with the Cdc28 (a cyclin dependent kinase). These complexes participate in actin polarization, which is crucial for hyphal growth (7, 30). On the other hand, low temperatures inhibit hyphal formation through the inhibitory activity of the molecular chaperone Hsp90 on the PKA signalling pathway (involving Ras1). Other environmental factors modulate filamentation through alternative signalling pathways, including pH (through the Rim101 pathway), hypoxia (involving transcription factors Efg1 and Efh1), low nitrogen (through transcription factor Cph1), and physical embedding of cells within a matrix (requiring transcription factor Czf1) (7, 30). Interestingly, farnesol blocks induction of hypha formation by most environmental signals, suggesting that it is involved in most of the filamentation signalling pathways (31, 32). Farnesol prevents hyphal initiation by downregulating the cAMP-PKA pathway (31, 32) and through an independent cAMP-PKA pathway (18).

Our data demonstrate that increased Leu-CUG incorporation in the *C. albicans* proteome triggers filamentation independently of environmental cues. In non-inducing conditions, high-level Leu-CUG translating strains showed robust filamentation (20% to 50% of germ tube formation) relative to wt cells. Also, both wt and T1, T1KO1, T2KO1 and T2KO2 cells induced >80% germ tubes at 37°C in medium containing serum. This is in line with previously reported data showing similar levels of filamentation in these inducing conditions (33) and proves that classical inducing mechanisms are not affected by Leu-CUG translation.

*C. albicans* proteins tolerate relatively well both Ser and Leu at CUG sites. For example, incorporation of 100% of Leu or Ser at the single CUG sites of the seryl-and leucy-tRNA synthetases (SerRS and LeuRS, respectively) did not affect tRNA aminoacylation, although the Leu isoforms were slightly more active than the Ser isoforms (23, 34). Also Leu-CUG isoforms of *Als3*, a cell wall adhesin of the core filamentation network (35) whose gene contains 2 CUGs, have slightly higher adherence to host substrates and flocculate in liquid medium, but no differences in protein stability were observed relative to the Ser-CUG isoforms (36). In another recent study, Leu-CUG translation decreased the stability of the protein kinase Cek1 without major structural alterations. Incorporation of Ser at this CUG site induced the autophosphorylation of the conserved tyrosine residue of the Cek1^231^ TEY^233^ motif and increased its intrinsic kinase activity *in vitro* (37). Cek1 is a key kinase of the MAPK cascade directly linked to morphogenesis in *C. albicans*. Therefore, Leu/Ser-CUG isoforms of *C. albicans* proteins are normally functional despite some differences in activity and/or specificity.

In this work, we established a mechanistic link between Leu-CUG translation, filamentation and the metabolite farnesol. This quorum sensing metabolite is released in the extracellular environment at high cell densities and inhibits hypha formation (19). In our non-inducing conditions, the robust filamentation induced by high level of Leu-CUG translation (20% to 50% of germ tube formation in T1, T1KO1, T2KO1 and T2KO2 strains) was accompanied by strong decrease in farnesol production (Fig. 2). 100 μM of exogenously added farnesol was sufficient to recover the filamentation phenotype in the strains that incorporated Leu at CUG sites at high level. (Fig. 3). A previous study showed that 300 μM of exogenously added farnesol affected the interaction of the mRNA with the small ribosomal subunit leading to translation inhibition to limit growth and filamentation (38), but the 100 μM of exogenously added farnesol used in this study is below this toxicity threshold.

Farnesol inhibits hyphal initiation through stabilization of the repressor of the hyphal transcriptional program (15, 18), but it is also important for the other phase of hyphal development, i.e., maintenance, and promotes the transition back to the yeast form (33). In this work, we did not investigate hyphal maintenance or hypha-to-yeast transition and it will be interesting to study these phenotypes in future works.

Similar to classical filament-inducing cues, Leu-CUG translation in T1 strain required degradation of the transcriptional repressor Nrg1 to activate filamentation (Fig.4). Indeed, high-level Leu-CUG translation strains showed decreased Nrg1 protein expression in non-inducing conditions. We also observed that this decrease was not a direct consequence of the Ser-for-Leu substitution at the CUG site of this protein (Fig. 5), rather an indirect consequence of the farnesol-sensing pathway, which is essential for hyphal initiation.

Downregulation of Nrg1 during hyphal initiation is achieved independently by the cAMP-PKA pathway and degradation triggered by its release from farnesol. In our study, while Nrg1 protein degradation was detected in high level Leu-CUG translating cells, no alterations in the cAMP-PKA transcriptional pathway were observed (Table 1). This may explain the lower levels of yeast-to-hypha transition (up to 45%) relative to the near complete filamentation observed in Δ*nrg1* mutants (15). Although statistically significant differential expression of hyphal-specific genes regulated by Nrg1, namely *ALS3* (39), *CEK1*, *ECE1* (6) and *HWP1* (15) was observed in T1 cells, the expression fold change was relatively small (<2). *ECE1* was the first hypha-specific gene identified and its transcript levels are related to cell elongation (40). It is a well-studied fungal peptide toxin critical for mucosal infection named candidalysin (41). *HWP1* encodes a mannoprotein with a C-terminal GPI anchor and is exclusively expressed in the hyphal cell surface. Hwp1 is a substrate of the mammalian transglutaminase (TGase) and regulates covalent attachments of germ tubes to host epithelial cells (42). Therefore, other factors may be involved in this phenotype, including the key players of the response to farnesol, namely the Cek1 MAPK pathway (43) or the Ras1-cAMP pathway (31).

Our data suggest that adaptive Leu-CUG translation expands the repertoire of factors that influence virulence traits, specifically yeast-to-hypha transition. It induces fungal morphogenesis at 30°C via a cAMP-PKA transcriptional regulation independent mechanism. Decreased farnesol levels upregulated the expression of the Sok1 kinase, which is essential for Nrg1 protein degradation.

How Leu-CUG translation alters the production of farnesol is an open question. Farnesol synthesis pathways are not fully understood in *C. albicans* yet, but this metabolite is produced by the ergosterol pathway through the dephosphorylation of farnesyl pyrophosphate (FPP). Three pyrophosphatases encoding genes are present in the *C. albicans* genome (*DPP1, DPP2*, *DPP3*), but only the role of Dpp3 in farnesol production has been experimentally established (44, 45). Dpp3 is not essential for farnesol synthesis (33) and previous works suggest that Dpp1 and Dpp2 also participate in FPP dephosphorylation (33, 46). Dpp1 and Dpp2 enzymes contain CUG-encoded residues, but only the Ser-for-Leu substitution in Dpp1 is predicted to destabilize the protein (Table S3). Our microarray data did not show any alteration in *DPP1*, *DPP2* and *DPP3* expression nor in the genes associated with the ergosterol biosynthesis pathway upstream of farnesol production (47) in the high-level Leu-CUG translating strain (T1) (Dataset S1). One possibility is that Leu incorporation at the CUG-site could destabilize Dpp1 and target it for degradation, contributing to farnesol decrease in conditions where Leu-CUG translation increases (Fig.6). This hypothesis will be tested in the near future.

**Fig 6.**
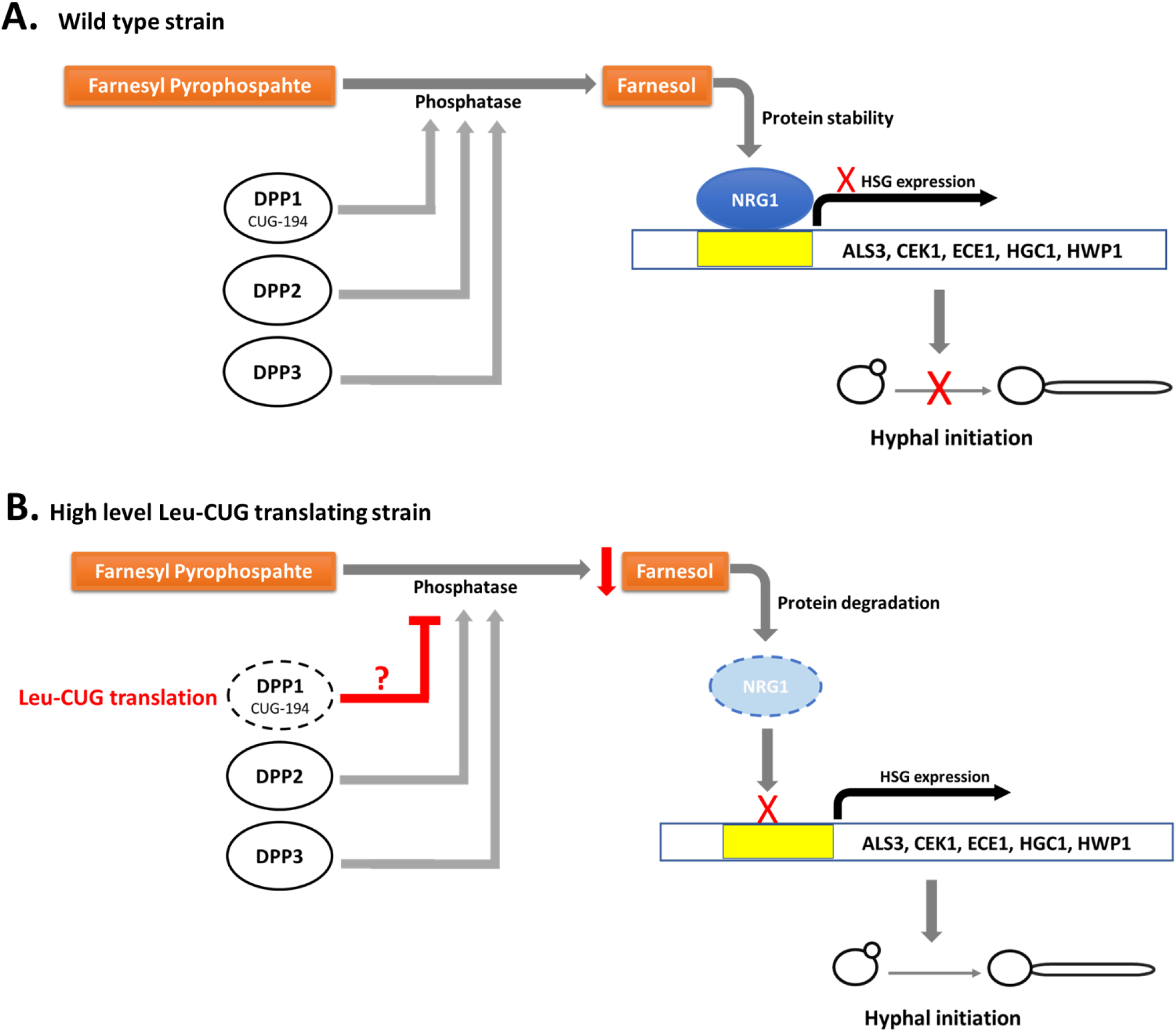
Schematic representation illustrating the modulation of filamentation by adaptive translation of the CUG codon in *C. albicans.* **A.** Regulation of hyphal initiation in wt *C. albicans* cells grown in non-inducing conditions, through the maintenance of Nrg1 protein stability. Presence of farnesol, produced from the dephosphorylation of farnesyl pyrophosphate catalyzed by Dpp3 and presumably by Dpp1 and Dpp2, prevents Nrg1 protein degradation. Nrg1 expression prevents hyphal development and expression of hypha-specific genes (e.g. *CEK1, ECE1, HWP1*). **B**. Proposed model for the regulation of filamentation in *C. albicans* high-level Leu-CUG translating cells in non-inducing conditions. Ser-for-Leu substitution at CUG-site could directly destabilize Dpp1 and target it for degradation, contributing to a decrease in farnesol production. The absence of farnesol leads to Nrg1 protein instability, promoting hyphal initiation and expression of hypha-specific genes.

## Material and methods

### Strains, media and growth conditions

*C. albicans* strains were routinely grown at 30°C in YPD (2% Bacto peptone, 2% dextrose, 1% yeast extract). Wild type and high level Leu-CUG translating strains are detailed in Table S1.

### GT assay

As a measure of filamentation, a germ tube (GT) assay was performed as described previously (16, 24) with minor modifications. Strains were grown overnight in liquid YPD at 30°C, pelleted, washed twice in PBS, resuspended in an equal volume of PBS, and diluted 1:100 into YPD 30°C, RPMI 37°C, RPMI+10% serum (Sigma-Aldrich) and YPD+Farnesol 100 μM (Sigma-Aldrich). Cultures were grown for 5 hours while shaking at 180 rpm. The morphology of cells was analyzed at the indicated timepoints using a Zeiss MC80 Axioplan 2 microscope. Germ tube formation was quantified from a minimum of 300 cells. A cell was counted as germ tube when an initial filament had formed (irrespective of hypha- or pseudohypha-like morphology). Each experiment was performed at least three times independently. Differences in percentages of GT formation were tested using Student’s t-test with the threshold of a *P* < 0.05 for statistical significance.

### Volatile metabolites determination using GC×GC–ToFMS

*C. albicans* strains were grown overnight in liquid YPD at 30°C, pelleted, washed twice in PBS, resuspended in an equal volume of PBS, and diluted 1:100 into YPD 30°C. Cultures were then grown for 5 hours while shaking at 180 rpm. Cell concentration was determined for three independent cultures of each strain using the TC20 automated cell counter (Bio-Rad). These results were used to normalize the total areas of each chemical feature detected, therefore allowing the determination of specific metabolite production per cell.

After incubation, 200 mL of each sample were collected and centrifuged at 10.000 rpm, at 4°C for 15 min. For headspace solid phase microextraction (HS-SPME), 20 mL (1/β ratio of 0.5) of supernatant were transferred into a 60 mL glass vial, via syringe with 0.20 μm filter pore. After the addition of 4g of NaCl (≥ 99.5%, Sigma-Aldrich) and stirring bar of 20×5 mm, the vial was capped with a silicone/polytetrafluoroethylene septum and an aluminium cap (Chromacol Ltd., Herts, UK). The samples were stored at −80°C until analysis.

The selected SPME device included a fused silica fiber coating, partially cross-linked with 50/30 μm divinylbenzene/carboxen™/polydimethylsiloxane StableFlex™ (1 cm), which comprehends a wide range capacity of sorbing compounds with different physicochemical properties (48). After defrost, vials were placed in a thermostated water bath so that headspace extraction was allowed to occur for 30 min, at 50°C ±0.1 and under continuous agitation at 350 rpm. Three independent aliquots were analysed for each condition under study.

The SPME fiber was manually introduced into the GC×GC–ToFMS injection port and exposed during 30 seconds for thermal desorption into heated inlet (250°C). The instrumental parameters were defined according to a previous metabolomics study (49). The inlet was lined with a 0.75 mm I.D. splitless glass liner and splitless injections mode were used (30 seconds). The LECO Pegasus 4D (LECO, St. Joseph, MI, USA) GC×GC– ToFMS system was comprised by an Agilent GC 7890A gas chromatograph (Agilent Technologies, Inc., Wilmington, DE), with a dual stage jet cryogenic modulator (licensed from Zoex) and a secondary oven, as well as mass spectrometer equipped with a ToF analyzer. An Equity-5 column (30 m × 0.32 mm I.D., 0.25 μm film thickness, Supelco, Inc., Bellefonte, PA, USA) and a DB-FFAP column (0.79 m x 0.25 mm I.D., 0.25 μm film thickness, J&W Scientific Inc., Folsom, CA, USA) were used for first (^1^D) and second (^2^D) dimensions, respectively. The carrier gas was helium at a constant flow rate of 2.50 mL min^−1^. The following temperature programs were used: the primary oven temperature was ranged from 40°C (1 min) to 140°C at 10°C min^−1^, and then to 200°C (1 min) at 7°C min^−1^. The secondary oven temperature program was 15°C offset above the primary oven. Both the MS transfer line and MS source temperatures were 250°C. The modulation period was 5 seconds, keeping the modulator at 20°C offset above primary oven, with hot and cold pulses by periods of 0.80 and 1.70 seconds, respectively. The ToF analyzer was operated at a spectrum storage rate of 100 spectra s^−1^, with mass spectrometer running in the EI mode at 70 eV and detector voltage of −1499 V, using an *m/z* range of 35-300. Total ion chromatograms were processed using the automated data processing software ChromaTOF^®^ (LECO) at signal-to-noise threshold of 200. For identification purposes, the mass spectrum and retention times (^1^D and ^2^D) of the analytes were compared with standards, when available. Also, the identification process was done by comparing the mass spectrum of each peak with existing ones in mass spectral libraries, which included an in-house library of standards and two commercial databases (Wiley 275 and US National Institute of Science and Technology (NIST) V. 2.0 - Mainlib and Replib). Moreover, a manual analysis of mass spectra was done, combining additional information like linear retention index (LRI) value, which was experimentally determined according to van den Dool and Kratz RI equation (50). A C_8_-C_20_ *n*-alkanes series was used for LRI determination (the solvent *n*-hexane was used as C_6_ standard), comparing these values with reported ones in existing literature for chromatographic columns similar to ^1^D column above mentioned. The majority (> 90%) of the identified compounds presented similarity matches > 800/1000. The Deconvoluted Total Ion Current GC×GC area data were used as an approach to estimate the relative content of each metabolite.

### GC×GC–ToFMS statistical analysis

Firstly, peak areas of all analytes were manually extracted from the chromatograms and used to build the full data matrix from *C. albicans* strains cultures. The complete list of these analytes is provided in Table S2, including the average areas of three independent assays for each strain under study. The significance of the analytes detected in the fungal cultures (GC areas normalized for the number of cells) were compared to the ones that were detected in the YPD medium (control), through a two-sided Mann-Whitney test (using the SPSS software 20.0 (IBM, New York, USA)).

Differences corresponding to *P* ≤ 0.05 were considered significant. Student’s t test was applied to this dataset to evaluate which metabolites were considered different between high-level Leu-CUG translating and control strains. The heatmap visualization was applied for this dataset using the MeV software (51).

### *In silico* prediction of the effect of single amino acid substitutions

PROVEAN (Protein Variation Effect Analyzer) (29) is an evolutionary trend-based classifier of amino acid substitutions. PROVEAN runs a Blast search and the clustering of BLAST hits is performed by CD-HIT with a parameter of 75% global sequence identity. The top 30 clusters of closely related sequences are used to generate the prediction. The prediction is based on the change, caused by a given variation, in the similarity of the query sequence to a set of its related protein sequences. For this prediction, the algorithm is required to compute a semi-global pairwise sequence alignment score between the query sequence and each of the related sequences (29).

### Western blot

Total protein fractions were analyzed under reducing conditions using 10% SDS-PAGE and blotted onto nitrocellulose membranes (Hybond ECL, Amersham) according to standard procedures. The His_6_-tag was detected using a mouse antibody against His tag (18184, Abcam, Cambridge, UK) at 1:5000 dilution. Bound antibody was visualized by incubating membranes with an IRDye680 labelled goat secondary antibody against mouse (Li-Cor Biosciences, Lincoln, NE, USA) at 1:10 000 dilution. Detection was performed using an Odyssey infrared imaging system (Li-Cor Biosciences). As an internal control a monoclonal anti-α-tubulin antibody (Sigma) was used.

### Genomic expression profiling

*C. albicans* strains were grown overnight in liquid YPD at 30°C, pelleted, washed twice in PBS, resuspended in an equal volume of PBS, and diluted 1:100 into YPD 30°C. Cultures were then grown for 5 hours while shaking at 180 rpm. 50 ml of exponentially growing cells were harvested and frozen overnight at −80°C. Total RNA was extracted using the hot-phenol method (52) with few modifications. Total RNA samples were treated with DNaseI (Amersham Biosciences) according to the commercial enzyme protocol and quantification and quality control was performed using the Agilent 2100 Bioanalyzer system. Gene expression profiling was performed using the Agilent protocol for One-Color Microarray Based Gene Expression Analysis Quick Amp Labeling v5.7 (Agilent Technologies). Briefly, cDNA was synthesized from 600ng of total RNA using AgilentT7 Promoter Primer andT7 RNA Polymerase Blend and labelled with Cyanine 3-CTP. Labelled cDNA was purified with RNeasy mini spin columns (QIAGEN) to remove residual Cyanine 3-CTP. Dye incorporation and quantification was monitored using the Nanodrop 1000Spectrophotometer. To prepare hybridization, 1.65μg of Cy3-labeled cRNA were mixed with the fragmentation mix (Blocking Agent and Fragmentation Buffer) and incubated for 30 min at 60°C. Finally, GEx Hybridization Buffer HI-RPM was added and the preparation was assembled in the custom-made Agilent arrays (GE1_105_Dec08). Slides were prepared using Agilent gasket slides according to the manufacturer instructions. Each hybridization was carried out for 17h at 65°C, in an Agilent hybridization oven. After washing and drying, the microarrays were scanned using the Agilent G2565AA microarray scanner (Agilent).

### Microarray data extraction and analysis

Probes signal values were extracted from microarray scan data using Agilent Feature Extraction Software (Agilent). Data were normalized using median centering of signal distribution with Biometric Research Branch BRB-Array toolsv3.4.0 software. Microarray data analysis was carried out with MEV software (TM4 Microarray Software Suite) (51). Student’s t test was applied to identify genes that showed statistically significant (*P* ˂0.05) differences in expression between control and high-level Leu-CUG translating strains.

## Data availability

The microarray raw data was submitted to the GEO database and has been given the following accession number: GSE164205.

## Supplemental material

Table S1: *Candida albicans* strains used in this study.

Table S2: Metabolites putatively identified in *Candida albicans* using GC×GC-ToFMS.

Table S3: PROVEAN prediction of the effect of Ser-for-Leu substitution on CUG-encoded residues of proteins.

Dataset 1: Complete list of gene expression in the high-level Leu-CUG translating T1 strain as revealed by transcript profiling.

Supplementary methods: Description of growth conditions, construction of plasmids and strains.

## Acknowledgements

This work was supported by FEDER (Fundo Europeu de Desenvolvimento Regional) funds through the COMPETE 2020, Operational Programme for Competitiveness and Internationalization (POCI), and by Portuguese national funds via Fundação para a Ciência e a Tecnologia, I.P. (FCT) under the projects PTDC/IMI-MIC/5350/2014, PTDC/BIA-MIB/31238/2017 and PTDC/BIA-MIC/31849/2017. The iBiMED and LAQV-REQUIMTE research units are supported by FCT funds under UIDP/04501/2020 and UIDB/50006/2020, respectively. CO is supported directly by a FCT grant (SFRH/BD/131637/2017). ARB is supported by national funds (OE), through FCT, I.P., in the scope of the framework contract foreseen in the numbers 4, 5, and 6 of the article 23, of the Decree-Law 57/2016, of August 29, changed by Law 57/2017, of July 19.

